# Motor memory under sleep deprivation: Can physical exercise support consolidation?

**DOI:** 10.64898/2026.01.07.698089

**Authors:** Guillaume Digonet, Thomas Lapole, Juliette Gelebart, Jérémie Bouvier, Mélanie Prudent, Vincent Pialoux, Maxime Pingon, Emeric Stauffer, Ursula Debarnot

## Abstract

Sleep deprivation (SD) is increasingly prevalent and known to impair declarative memory, yet its impacts on the acquisition and consolidation of procedural skills remains unknown. Physical exercise has emerged as a promising intervention for promoting learning and plasticity, but its potential to mitigate SD-induced deficits has never been tested. Here, we investigated whether SD disrupts sequential motor learning (SML) and whether high-intensity interval exercise (HIIE) performed after acquisition can influence consolidation. Forty-eight participants were randomly assigned to one of four groups combining one night of SD or normal sleep condition with HIIE or control intervention. SD elevated cortisol and sleepiness without affecting brain-derived neurotrophic factor or corticospinal excitability. Behaviorally, SD selectively impaired the movement time execution of SML acquisition, while HIIE reduced the movement accuracy during consolidation, regardless of sleep condition. These findings reveal a component-specific vulnerability of procedural memory to SD and challenge the assumption that exercise consistently enhances consolidation following motor learning.

## Introduction

Sleep is a biologically active state essential for the regulation of memory across multiple levels of brain function^1,2^. From synaptic plasticity to large-scale cognitive operations, sleep facilitates the encoding, consolidation, and retrieval of information, particularly within hippocampus-dependent systems underlying declarative and skilled motor memory^3,4^. Although an extensive body of literature has established the critical role of sleep in memory, the growing prevalence of sleep deprivation (SD) underlines the urgent need to further elucidate the neurobiological mechanisms supporting sleep-dependent memory processing^5,6^.

Several evidence from human studies has established that SD leads to significant impairments in hippocampus-dependent declarative memory systems^6^. Notably, memory encoding appears particularly vulnerable to sleep loss, with persistent impairments observed following total SD, whereas retrieval mechanisms remain relatively intact^5^. At the neurobiological level, SD has been associated with increased cortisol levels^7^, whereas its cognitive deficits have been linked to a downregulation of brain-derived neurotrophic factor (BDNF)^8^, a key neurotrophin involved in neuroplasticity and memory formation^9^. Although the effects of SD on declarative memory are increasingly well understood, its impact on skilled motor memory, which also relies on hippocampal function^10^, remains largely unexplored. Existing evidence in the deleterious effect of SD in procedural memory has focused exclusively on the consolidation^11^, leaving a critical gap in our understanding of how sleep loss affects the initial encoding of motor skills. To date, only the study by Fischer et al. (2002)^12^ has indirectly addressed this question, including an underpowered SD group that was not central to the original design and reported no effect of SD on sequential motor learning (SML). Since then, the impact of sleep loss on both motor skill acquisition and consolidation has remained an open and unresolved question.

Physical exercise has recently gained attention as a potential strategy to mitigate the adverse effects of SD in declarative memory^13,14^. Evidence from animal studies supports this view, demonstrating that physical exercise can prevent SD-induced disruptions in neuroplasticity and learning^15^. In human studies, exercise performed shortly after learning has been shown to enhance motor memory consolidation, as evidence 24 hours later ^16,17^. These gains are most consistently reported following high-intensity interval exercise (HIIE), which induces lactate production and BDNF release^18^. Together, lactate provides the metabolic support required for consolidation^19^, while BDNF opens a time-sensitive window for synaptic plasticity^20^, thereby fostering a neurochemical state that favors motor memory improvements over 24 hours and up to 7 days^21^. Moreover, HIIE has emerged as the most effective modality to modulate corticospinal excitability^22^, where elevations in motor-evoked potentials (MEP) and reductions in short-interval intracortical inhibition (SICI) serve as predictors of behavioral performance gains during consolidation^23,24^. Altogether, these findings suggest that HIIE promotes motor memory consolidation and may offer a potential means to counteract memory impairments induced by SD.

This study investigated whether SD alters the acquisition of SML, and whether HIIE performed after learning can support memory consolidation under both rested and sleep-deprived conditions. We further examined biological (cortisol, BDNF, lactate) and neurophysiological (MEP, SICI) markers of plasticity, hypothesizing that SD would increase cortisol and decrease BDNF levels, whereas HIIE would elevate lactate and BDNF, enhance MEP, and reduce intracortical inhibition.

## Materials and Methods

### Participants

Forty-eight healthy right-handed participants (25 females; aged 18-35), confirmed by the Edinburg Handedness Inventory (score > 0.4)^25^, with good sleep quality (PSQI score < 7) ^26^, were randomly assigned to parallel group SLEEP_HIIE_, SLEEP_REST_, SD_HIIE_ or SD_REST_ group based on the night (SD or SLEEP) prior motor acquisition and the physical intervention (HIIE or REST) post-acquisition (Fig. 1). Participants were pseudorandomized with online case report form based on their gender to ensure equal distribution across the four groups. Exclusion criteria included (1) ferromagnetic implants, (2) history of neurosurgery, (3) seizures, major head trauma, cardiovascular or respiratory disease, alcoholism, or drug addiction, (4) recent limbs injuries (< 6 months), and (5) any psychiatric or neurological disorders, or a resting heart rate exceeding 100 bpm. Musicians with more than five years of practice were excluded due to advanced finger dexterity, as well as participants scoring below four on the Corsi test assessing visuospatial memory span (PsyToolkit ; See table 1 for participants information)^27^. Participants were instructed to abstain from alcohol and caffeine for 24h prior to and during the experimental sessions. The present research was conducted in accordance with ethical guidelines and approved by the ethical committee of Île de France X (2023-A00155-40). It complies with the Declaration of Helsinki and was preregistered on ClinicalTrials.gov (NCT05910814). The recruitment period for this study started on 16/06/2023 and ended on 20/09/2024. Written informed consent was obtained from all participants after a full explanation of the procedures. Participants received a compensation fee for their participation.

**Figure 1:**
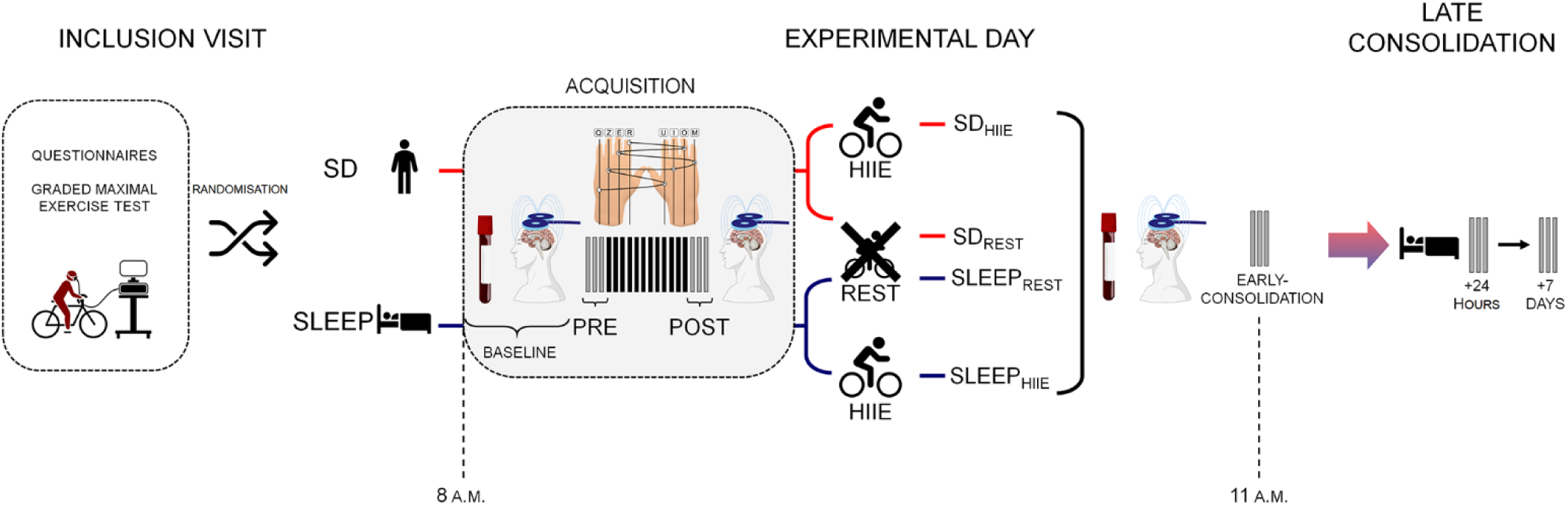
Experimental design overview. Following the inclusion visit, which involved questionnaires and a graded maximal exercise test (for HIIE groups only), participants were randomly assigned to one of four experimental groups: SD_HIIE_, SD_REST,_ SLEEP_REST_, SLEEP_HIIE_. The night before the experimental day, participants either experienced total sleep deprivation (SD) or had a full night of sleep (SLEEP). On the experimental day, blood samples were collected before (Baseline) and after SML acquisition to measure cortisol and BDNF levels, and TMS assessments (MEPs, SICI) were performed at the same point. Participants performed a bimanual 8-item SML across 12 blocks of practice, and performance (i.e. accuracy and movement time variables) was evaluated before and after the acquisition phase (PRE and POST). Following post-learning TMS assessment, participants completed either high-intensity interval exercise (HIIE) or resting intervention (REST). Consolidation performance was probed immediately after (Early), and at 24 hours and 7 days (Late).

**Table 1:** Characteristics of the participants for each group.

| | $SLEEP_{REST}$ | $SLEEP_{HIIE}$ | $SD_{REST}$ | $SD_{HIIE}$ | P value |
| --- | --- | --- | --- | --- | --- |
| AGE | $23.3 \pm 3.1$ | $24.7 \pm 3.9$ | $24.8 \pm 2.7$ | $23.7 \pm 1.6$ | 0.28 |
| VO2max (ml/min/kg) | | $43.8 \pm 8.1$ | | $45.5 \pm 6.1$ | 0.49 |
| MAP (Watt) | | $227.5 \pm 51.8$ | | $233.8 \pm 57.1$ | 0.93 |
| GPAQ | $4031.7 \pm 2721.7$ | $2551.7 \pm 1463.3$ | $2435.0 \pm 934.0$ | $3328.3 \pm 2328.3$ | 0.45 |
| EDINBURGH | $0.9 \pm 0.16$ | $0.85 \pm 0.12$ | $0.85 \pm 0.15$ | $0.85 \pm 0.17$ | 0.70 |
| CIRCADIAN TYPOLOGY | $56.5 \pm 7.0$ | $52.4 \pm 7.4$ | $55.5 \pm 4.0$ | $53.4 \pm 5.8$ | 0.60 |
| PSQI | $4.5 \pm 2.6$ | $4.75 \pm 2.9$ | $4.9 \pm 2.8$ | $4.2 \pm 1.9$ | 0.89 |
| CORSI | $6.1 \pm 0.7$ | $6.3 \pm 0.8$ | $6.3 \pm 1.0$ | $6.3 \pm 1.0$ | 0.91 |

P-values correspond to the results of Kruskal Wallis or Wilcoxon tests. GPAQ = global physical activity questionnaire; MAP = maximal aerobic power; PSQI = Pittsburgh Sleep Questionnaire Index.

The sample size was determined using G*Power software (version 3.1.9.7)^28^ based on a priori power analysis aimed at detecting an effect size *f* of 0.25 (medium effect), based on the meta-analysis from Newburry et al. (2021)^6^, which examined the effect of SD on declarative memory and reported a medium to large effect. The statistical power and alpha were set at 0.8 and 0.05, respectively, yielding an estimated sample size of 48 participants (i.e. 12 participants in each group).

### Design procedure

At inclusion visit, participants completed the Pittsburgh questionnaire^26^, Corsi test^27^, global physical activity questionnaire (GPAQ)^29^, circadian typology (30 > Score < 70)^30^ and Edinburgh handedness inventory^25^. The participants enrolled in HIIE groups underwent a graded maximal test to assess their VO2max and their maximal aerobic power (MAP).

Two days before the experiment, participants wore an actimeter (wGT3X-BT, Pensacola, USA), until the morning of the experimental day. SD participants arrived at the laboratory at 10:00 p.m. and were continuously monitored by the experimenter to ensure complete SD. SLEEP participants slept at home under usual conditions. The experimental day began at 8:00 a.m. at the Croix-Rousse Hospital (Lyon, France) with blood sampling and TMS measurements. SLEEP participants self-reported a sleep duration of 07:28 min (± 1:14), whereas SD participants had been awake for 24h (± 1:15). Performance at the bimanual 8-element SML task was then assessed across three blocks of PRE-ACQUISION test before engaging in the practice phase lasting 12-blocks. Performance was reassessed on three other blocks at POST-ACQUISITION test and, a second neurophysiological TMS assessment was conducted. Participants then either performed a 17-min HIIE on a cycle ergometer (Ergoline GmbH, Germany) or remained seated watching a documentary (REST). The HIIE protocol included a 2-minute warm-up (50 W), followed by three rounds of 3-min bouts at 80 % of maximal aerobic power, each interspersed by 2-min of active recovery at 40 %. The session ended with a 3-min cooldown at 50 W. Lactate samples were collected before and 2-min after HIIE or REST, and a second blood sample was collected 5-min post-intervention. This was immediately followed by a third TMS assessment. To examine the effect of HIIE and REST on EARLY-CONSOLIDATION, participants were then re-evaluated on three blocks of SML task. Finally, performance was assessed remotely at 24 Hours and 7 Days later to evaluate late consolidation as illustrated Fig. 1.

### Sequential motor learning task

Participants were seated on a chair positioned 50 cm in front of a computer screen. The fingers were placed on the alphabetical AZERTY keyboard, with Q corresponding to the left little finger, Z to the left ring finger, E to the left middle finger, R to the left index, U to the right index, I to the right middle finger, O to the right ring finger, M corresponding to the right little finger and finally, both thumbs were placed on the space bar. The sequence of eight movements required pressing the keys in the following order R-O-E-M-I-Z-U-Q, and systematically validating the sequence by pressing the space bar with the right thumb (i.e. the left thumb was not used). Participants were instructed not to correct conscious errors but instead to press the space bar to start a new sequence. Each experimental block required participants to repeat the sequence eight times as accurately and as quickly as possible. Each block began with the message “Are you ready?” on the screen, followed by a 3-s countdown before the first sequence, and finished with a 20-s rest period. The sequence remained displayed on the screen during blocks and rest periods. The task was performed using Psytoolkit. The order and the timing of each keypress was recorded and used to calculate the ACCURACY (i.e. the percentage of correct sequences achieved within a block), with a maximum of 8 correct sequences per block, and the MOVEMENT TIME (i.e. average time, in milliseconds, to complete a correct sequence in a block).

### Graded maximal test

The graded maximal exercise performed by participants from HIIE groups was conducted on a cycle ergometer (Ergoline GmbH, Allemagne). The test began with a 3-min warm-up at a power output of define the subject’s physical condition. After the warm-up period, participants proceeded directly to the incremental ramp protocol, consisting in a 15 or 25 W increase each minute, depending on the subject’s physical condition, until exhaustion was reached, and was used to assess maximal aerobic power and corresponding VO2max (Quark CPET system, Cosmed, Italy). The maximal power output achieved during this test was used to adjust the intensity of HIIE.

### Neurophysiological assessment

Surface electromyography (EMG) was recorded from the right first dorsal interosseous (FDI) muscle using Ag/AgCl electrodes (Kendall™ Medi-Trace®) in a monopolar configuration. EMG signals were amplified (ML138, ADInstruments), digitized at 2 kHz (PowerLab 16/30-ML880/P, ADIstruments, Bella Vista, Australia) and bandpass filtered online between 20 and 500 Hz. Data were analyzed ousing Labchart 8 software. MEPs and SICI were elicited using simple or paired pulse TMS delivered over the left M1 via a figure-of-eight coil (70-mm, Magstim 200, UK). The coil was positioned at a 45° angle to the sagittal plane to induce a postero-anterior current and marked on a cap to ensure consistent placement for each TMS measurement. The individual resting motor threshold (rMT) was defined as the lowest intensity that evoking at least three MEPs > 0.05 mV out of five. MEPs were elicited using a single magnetic pulse at 120 % of the rMT, while SICI were evoked by paired-pulse stimulation with a conditioning stimulation at 70 % of rMT and a test stimulation at 120 %, separated by a 3-ms interval ^31^. During each TMS session, 10 MEPs and 10 SICI responses were recorded; the mean peak-to-peak amplitudes at each time point (PRE- and POST-ACQUISITION, EARLY-CONSOLIDATION) were used for analysis. MEPs amplitude was expressed as a percentage of maximal M-wave (Mmax), and SICI was quantified as the mean ratio between conditioned responses amplitude and MEPs amplitude. To account for changes in sarcolemmal excitability, MEPs were normalized to Mmax, obtained via supramaximal electrical stimulation of the right ulnar nerve (DS7R, Digitimer) using a bipolar felt pad electrode (30 mm, E.SB020/4mm). A single 1-ms rectangular pulse (max 400 V) was applied, starting at 5 mA and increased in 2-mA steps until Mmax was reached.

### Biological and sleepiness assessment

Biological assessment included plasma measurements of BDNF, cortisol and lactate levels. For BDNF and cortisol, venous blood samples (5 ml) were collected from the antecubital vein after the experimental night, either following SD or normal sleep (8:00 a.m.), and after the intervention, either HIIE or REST. Samples were centrifuged at 3000 x g for 10 min. The plasma was separated from the serum and was kept at -80°C until analysis. BDNF concentrations and cortisol were measured using enzyme-linked immunosorbent assay (ELISA) Kit (MyBioSource, San Diego, U.S.A) according to the manufacturer’s instructions. Capillary blood lactate was collected from the fingertip before and after HIIE or REST intervention and analyzed using a lactate analyzer device (Lactate Pro 2, Arkay, Kyoto, Japan). At the beginning of the experimental day, participants completed a Psychomotor Vigilance Test (PVT)^32^, consisting of a 2-minute reaction time task. Subjective wakefulness was assessed before each test session using the Stanford sleepiness scale (SSS)^33^.

### Statistical analysis

All the following analyses were performed on the R software (v.4.4.1) with the packages {lme4}, {glmmTMB} {emmean}, and {lmer}. Outcomes of the models (F-value, degrees of freedom, effect size) were reported only for significant factors, while p-values were reported for both significant and non-significant factors. All information for each factor or interaction can be found in Table S1 in the supplementary data. Baseline analyses were conducted using Wilcoxon tests to assess the influence of SD on MEPs, SICI, Cortisol, BDNF and Vigilance prior to task acquisition. SSS was examined though linear mixed model with fixed effects of NIGHT, INTERVENTION and TIME. To test the effects of SD on acquisition stage for MOVEMENT TIME and ACCURACY variable, a generalized linear mixed model (GLMM) with fixed effects of NIGHT (SD, SLEEP) and TIME (PRE-ACQUISITION, POST-ACQUISITION), and subject-specific random effects were computed. A linear model was further used to compare PRE- to POST-ACQUISITION changes using this formula [((POST-ACQUSITION – PRE-ACQUISITION) / PRE-ACQUSITION) × 100] in MOVEMENT TIME and ACCURACY between SLEEP and SD. MEP and SICI were analyzed using GLMM using the fixed effect NIGHT and TIME (PRE-ACQUISITION, POST-ACQUISITION). To analyze the consolidation effects the same GLMM for MOVEMENT TIME and ACCURACY were computed but using the fixed effects NIGHT (SD, SLEEP), TIME (POST-ACQUISITION, EARLY-CONSOLIDATION, 24 Hours, 7 Days) and INTERVENTION (HIIE, REST) and subject-specific random effects. While MEP and SICI were analyzed using the fixed effects NIGHT, TIME (POST-ACQUISITION, EARLY-CONSOLIDATION) and INTERVENTION. Blood ΔBDNF, Δcortisol and Δlactate were calculated as the difference between the second and first samples (i.e. Second sample – First sample). Δlactate, Δcortisol and ΔBDNF were analyzed using linear model with fixed effects of NIGHT and INTERVENTION.

## Results

### Physiological and sleepiness state following sleep deprivation or sleep

No differences were found between the SD and SLEEP groups in baseline BDNF levels (p = 0.222; Fig. 2A), MEP (p = 0.814) or SICI (p = 0.809). However, participants in the SD group showed higher baseline cortisol levels (W = 364; p = 0.004; r = 0.421; Fig. 2B) and reduced vigilance, as reflected by slower reaction times on a psychomotor vigilance task, compared to the SLEEP groups (W = 369.5; p = 0.022; r = 0.340; Fig. 2C). Analysis of Stanford sleepiness scale (SSS) scores revealed a significant TIME*NIGHT interaction (F(4,176) = 46.371; p < 0.001; 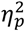 = 0.513; Fig. 2D), while all other interactions were nonsignificant (all p > 0.28). Post-hoc comparisons indicated that subjective sleepiness was higher in the SD than in SLEEP groups at PRE, POST and EARLY-CONSOLIDATION (all p < 0.001), while not different at 24 Hours (p = 0.208), and lower at 7 Days (p = 0.017). Moreover, a marked reduction in sleepiness was observed in SD groups between late consolidation (24 Hours and 7 Days) compared to PRE, POST, and EARLY-CONSOLIDATION (p < 0.001).

**Figure 2:**
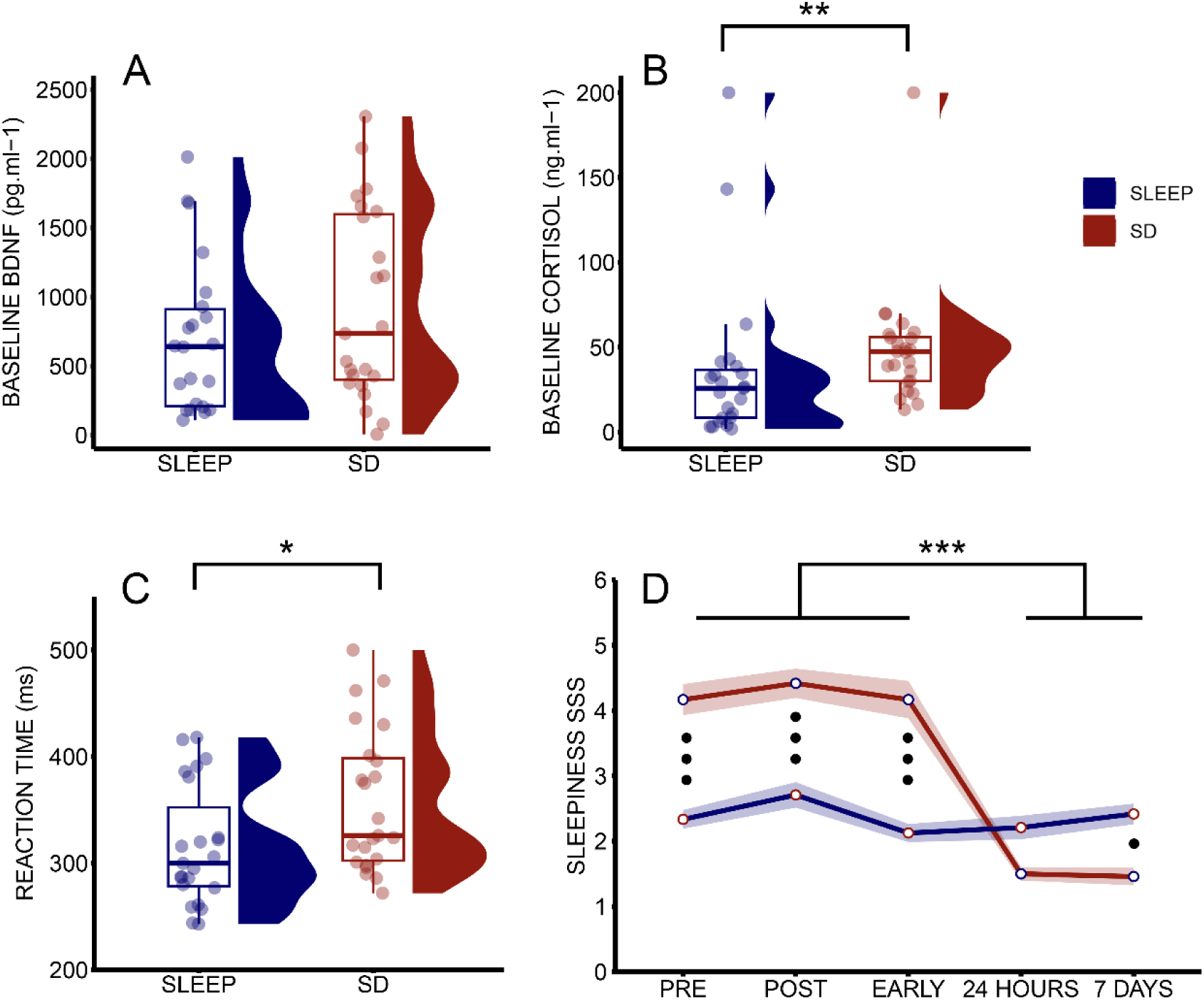
Baseline levels of BDNF, cortisol and vigilance after a night of SLEEP or sleep deprivation (SD). **A)** Baseline BDNF levels were equivalent between SLEEP and SD groups (p = .22). **B)** Baseline cortisol levels were higher in SD compared to SLEEP condition (p = .004). **C)** Objective vigilance assessed at the beginning of the experimental day using a psychomotor vigilance task was slower for SD compared to SLEEP group (p = .022). **D)** Temporal dynamics of subjective sleepiness across sleep conditions and time of testing. Red and blue lines represent SD and SLEEP groups respectively; shade areas indicate SEM. Higher scores reflect greater subjective sleepiness (i.e. lower vigilance). SD groups reported higher sleepiness than SLEEP groups at PRE-, POST-ACQUISITION, and EARLY-CONSOLIDATION (••• p < .001), but not at 24 Hours; at 7 Days, the SD group reported lower sleepiness (• p < .05). A significant drop in sleepiness was observed in the SD group at 24 Hours compared to all prior timepoints. * p < .05; ** p < .01; *** p < .001.

### Acquisition of motor movement time, but not accuracy, is affected by sleep deprivation

The analysis of MOVEMENT TIME (i.e. mean total time required to execute each correct sequence), revealed a main effect of TIME (χ²(1) = 331.44; p < 0.001), no main effect of NIGHT (p = 0.536), and a TIME*NIGHT interaction (χ²(1) = 5.873; p = 0.015; Fig. 3A). Post-hoc tests confirmed an improvement from PRE- to POST-ACQUISITION in both the SD and SLEEP groups (p < 0.001), however, percentage gains were attenuated following SD, with smaller gains in MOVEMENT TIME (+38.9 % ± 11.2) compared to the SLEEP groups (+46.8 % ± 12.2; W = 170; p = 0.015; r = 0.351; Fig. 3B). Moreover, movement time improvements during acquisition correlated negatively with subjective sleepiness (r(46) = -0.341, p = 0.018; Fig S1). In contrast, ACCURACY (i.e. percent of achievement in the number of correct sequences within a block, maximum of 8 sequences per block) showed no main effects of TIME (p = 0.510) or NIGHT (p = 0.417), and no TIME*NIGHT interaction (p = 0.147; Fig. 3C), in line with the reported percentage changes in ACCURACY (SLEEP: +3.1 % ± 24.4; SD: +4.3 % ± 25.9; p = 0.687; Fig. 3D).

**Figure 3:**
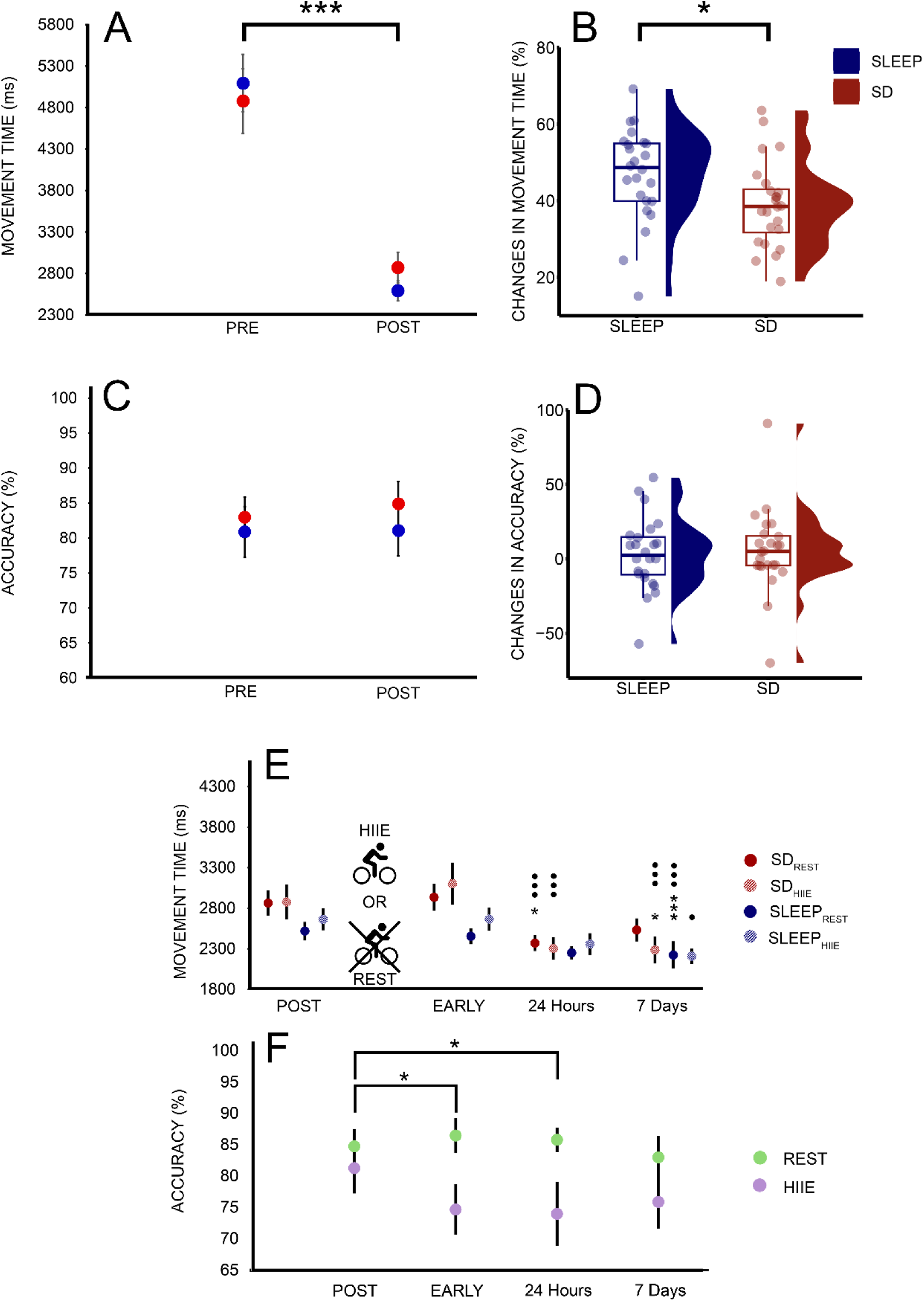
Effects of sleep condition on MOVEMENT TIME and ACCURACY during SML acquisition and consolidation. **A)** MOVEMENT TIME (ms) improved from PRE- to POST-ACQUISITION in both SLEEP (blue) and SD (red) (*** p < .001). **B)** Percentage change in MOVEMENT TIME showed greater performance gains in the SLEEP groups (+46.8 % ± 12.2) compared to the SD groups (+38.9 % ± 11.2). **C)** No change in ACCURACY from PRE- to POST-ACQUISITION in either group. **D)** Percentage change in ACCURACY did not differ between sleep conditions (SLEEP: +3.1 % ± 24.4; SD: +4.3 % ± 25.9). **E)** MOVEMENT TIME at POST-ACQUISITION, EARLY, 24 Hours and 7 Days. SD groups are shown in red, SLEEP groups in blue; REST conditions are shown as filled circles, HIIE as hatched circles. While no improvements were evident in EARLY-CONSOLIDATION, all groups displayed faster performance at later stages. * indicates significant differences from POST-ACQUISITION (* p < .05; *** p < .001), and • indicates differences from EARLY-CONSOLIDATION (• p < .05; ••• p < .001). **F)** ACCURACY at the same timepoints, by intervention type (green: REST; purple: HIIE). Accuracy significantly declined in the HIIE groups during EARLY and 24 Hours consolidation, but remained stable in REST. * p < .05. Errors bars represent SEM.

### High-intensity interval exercise impairs accuracy during motor memory consolidation

Analysis of HIIE vs REST interventions revealed a main effect of TIME for the MOVEMENT TIME (χ²(3) = 111.386; p < 0.001), with no main effect of NIGHT (p = 0.417) or INTERVENTION (p = 0.934). While no two-way interactions reached significance (all > 0.06), a three-way interaction between TIME*NIGHT*INTERVENTION emerged (χ²(3) = 7.981; p = 0.046; Fig. 3E). Post-hoc analyses showed a stabilization of MOVEMENT TIME between POST-ACQUISITION and EARLY-CONSOLIDATION in any group (p > 0.9 for each), with consolidation-related gains only emerging thereafter in group-specific patterns. In the SD_HIIE_ group, MOVEMENT TIME improved at 24 Hours and 7 Days relative to EARLY-CONSOLIDATION (both p < 0.001), and at 7 Days compared to POST-ACQUISITION (p = 0.012). In the SD_REST_ group, faster performance was found at 24 Hours compared to both EARLY (p = 0.001) and POST-ACQUISITION (p = 0.015). Similar improvements were observed at 7 days in both SLEEP_HIIE_ (vs EARLY; p = 0.029) and SLEEP_REST_ groups (vs. POST and EARLY; both p < 0.001). No difference in percentage change in MOVEMENT TIME were found (see Supplementary Data).

In contrast, ACCURACY was affected by a TIME*INTERVENTION interaction (χ²(3) = 10.712; p = 0.013), despite no main effects of NIGHT (p = 0.479), INTERVENTION (p = 0.092), nor TIME*NIGHT (p = 0.83), NIGHT*INTERVENTION (p = 0.248), and TIME*NIGHT*INTERVENTION interactions (p = 0.428). Post-hoc analysis revealed that ACCURACY declined in the HIIE groups at EARLY (p = 0.020) and 24 Hours (p = 0.045) relative to POST-ACQUISITION (Fig. 3F), whereas no such decline was observed in the REST groups. Percentage changes over time were not significant (see Supplementary Data).

### Biomarker modulation by sleep and exercise

ΔCORTISOL responses revealed a NIGHT*INTERVENTION interaction (F(1,41) = 10.927; p = 0.002; 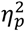 = 0.210), with greater decrease observed in the SD_HIIE_ compared to SLEEP_HIIE,_ and SD_REST_ (both p < 0.05; Fig. 4A), but not as compared to SLEEP_REST_ (p = 0.498). HIIE significantly increased ΔBDNF concentrations compared to REST (F(1,40) = 6.137; p = 0.017; 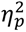 = 0.133; Fig. 4B), independently of sleep condition (NIGHT: p = 0.167; NIGHT*INTERVENTION: p = 0.596). Similarly, ΔLACTATE levels were markedly elevated following HIIE relative to REST (F(1,40) = 493.03; p < 0.001; 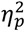 = 0.583; Fig. 4C), with no influence of sleep condition or interaction effects (both p > 0.9).

**Figure 4:**
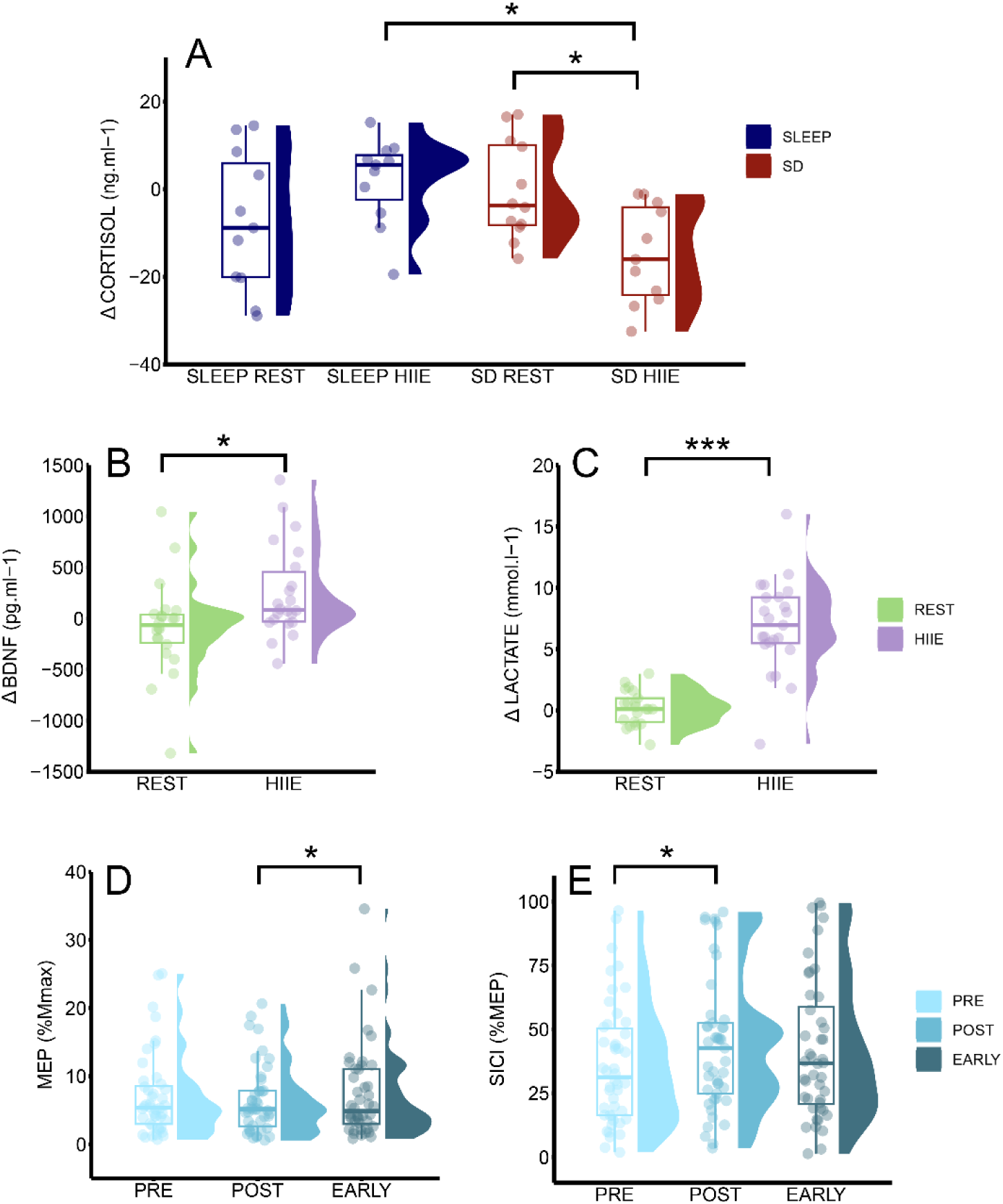
Biological marker following HIIE and REST interventions and neurophysiological responses during acquisition and early-consolidation. **A)** Interaction between sleep and intervention was observed for cortisol. SD_HIIE_ led to larger cortisol decrease compared to SD_REST_ and SLEEP_HIIE_. **B)** HIIE induced a greater increase in BDNF compared to REST, irrespective of sleep condition. **C)** Lactate levels increased after HIIE relative to REST. **D)** Corticospinal excitability (MEP) increased from POST-ACQUISITION to EARLY-CONSOLIDATION, regardless of sleep condition and intervention. **E)** SICI increased (i.e. reduction of intracortical inhibition) from PRE- to POST-ACQUISITION across all groups. Squares indicate SD participants; circles indicate SLEEP participants. Boxes show median and interquartile range. * p < .05; *** p < .001.

### Plasticity-related changes following motor learning and physical exercise intervention

After SML acquisition, analysis of MEP revealed no main effect of TIME (p = 0.465), NIGHT (p = 0.815) or TIME*NIGHT interaction (p = 0.198). In contrast, SICI showed no main effect of NIGHT (p = 0.675), and no TIME*NIGHT interaction (p = 0.935), but a main effect of TIME (χ²(1) = 5.867; p = 0.015) with reduced inhibition from PRE to POST-ACQUISITION (p = 0.015; Fig. 4D). Following the intervention, MEP exhibited a main effect of TIME (χ²(1) = 4.598; p = 0.032) with higher MEP amplitudes at EARLY-CONSOLIDATION compared to POST-ACQUISITION (p = 0.025; Fig. 4E). No effects of NIGHT (p = 0.466), INTERVENTION (p = 0.236), or interactions were detected (all p > 0.18). For SICI, a main effect of INTERVENTION was observed (χ²(1) = 3.98; p = 0.046), with lower inhibition in REST compared to HIIE groups (p = 0.048). No significant effects of analysis revealed no main effects of TIME (p = 0.550) or NIGHT (p = 0.925), or interaction terms were observed (all p > 0.24).

## Discussion

This study examine the effects of a full night of SD on SML, while also evaluating whether HIIE can promote memory consolidation under both SD and SLEEP conditions. To elucidate underlying mechanisms, we assessed vigilance, biological markers, and corticospinal excitability. Participants in the SD groups began the experiment with reduced vigilance and elevated cortisol relative to SLEEP groups, whereas BDNF levels and TMS-related measures remained comparable. Main findings revealed that SD selectively impaired the movement time, but not the accuracy, of SML acquisition. An intracortical disinhibition occurred following acquisition regardless of sleep status, suggesting that neuroplasticity-related processes were preserved despite SD. Importantly, although HIIE increased both BDNF and lactate levels, it affected accuracy during consolidation irrespective of the sleep status.

### Neurophysiological and attentional state following sleep deprivation or normal sleep

At the start of the experimental day, no differences in MEP, SICI and BDNF were observed between SLEEP and SD conditions, suggesting that one night of total SD does not impair baseline neuroplasticity markers. These results align with studies reporting no change in motor system excitability after SD^34^, although others have documented modulations^35,36^. Our initial hypothesis of reduced BDNF levels after SD, based on Kuhn et al. (2016)^8^, was not supported, with stable levels observed in line with Sochal et al. (2024)^37^ and Gorgulu et Caliyurt (2009)^38^, while other study have instead reported increases^39^. Discrepancies between studies likely reflect methodological and individual differences, including SD duration (e.g. 24h vs. 36h), the biological matrix used (serum vs plasma), participants’ fitness levels, or BDNF polymorphisms^37^. In contrast, cortisol levels were markedly elevated following SD, supporting its role as a physiological stressor and reflecting the disruption of slow-wave sleep-dependent mechanisms that normally suppress cortisol release and maintain circadian homeostasis^7,40^. SD also led to slower reaction times and increased subjective sleepiness, replicating classic signatures of decreased alertness after sleep loss^41,42^. Together, these findings suggest that although one night of SD alters stress-and vigilance-related markers, the neurophysiological readiness for plasticity remains intact.

### Acquisition of motor movement time, but not accuracy, is affected by sleep deprivation

One key finding is that SD selectively reduced movement time gains during SML acquisition without affecting accuracy. Neurophysiologically, we observed intracortical disinhibition, a pattern typically associated with motor learning under rested conditions, and thought to reflect modulation of GABAergic inhibitory mechanisms^44^ a crucial process in SML^45^. The preservation of this neurophysiological pattern, together with stable BDNF levels, suggests that plasticity mechanisms remained functional, and that the attenuated movement time improvements under SD likely reflects reduced motor execution efficiency rather than impaired learning processes. This interpretation is consistent with the dual-process model proposed by Hikosaka et al. (2002)^46^, which distinguishes between goal-based and movement-based components of SML, each supported by distinct neural substrates. The goal-based component involves the spatial representation of the sequence, relying on explicit processing, prefrontal working memory and hippocampal binding to integrate discrete elements into a coherent spatiotemporal structure^47^. In contrast, the movement-based component reflects implicit motor execution and progressive automatization, typically marked by movement time gains and underpinned by the supplementary motor area, striatum and cerebellum^46^. In our study, the preservation of accuracy despite SD suggests that spatial encoding and its associated prefrontal–hippocampal mechanisms remained intact. Conversely, the attenuated movement time improvements observed in the SD groups point to a disruption of movement-based processes, possibly reflecting reduced motor readiness or execution efficiency, in line with striatal alterations previously reported under SD^48^. As a complementary mechanism, reduced vigilance may have contributed to the impaired motor efficiency observed under SD. Slower reaction times, elevated subjective sleepiness, and the negative correlation between sleepiness and movement time execution indicate that diminished alertness and sustained attention may have further constrained movement time performance during the acquisition phase. Together, these findings highlight a dissociation between preserved hippocampal-dependent spatial learning and disrupted motor execution under SD, likely reflecting striatal vulnerability with reduced cognitive arousal. While this study focused on acute total SD, real-world sleep disruptions more commonly involve chronic partial restriction, often due to insomnia, delayed sleep onset, or fragmented sleep ^49^. Future research should therefore investigate partial sleep loss, as in the declarative memory study by Frimpong et al. (2023)^13^, since this experimental design may better capture ecologically relevant conditions and their impact on procedural memory.

### High-intensity interval exercise impairs accuracy during motor memory consolidation

A second key finding was that accuracy declined following HIIE during early-consolidation remained impaired at 24 hours, regardless of sleep condition, despite expected elevations in BDNF and lactate. This effect contrasts with previous studies showing improved motor performance in visuomotor adaptation following HIIE, with benefits typically emerging at 24 hours^17,50^. The discrepancy may reflect the specificity of explicit SML, where the accuracy component was selectively impaired, while execution movement time remained stable. This pattern suggests that HIIE disrupted goal-based components of SML, which rely on the functional interaction between the prefrontal cortex and hippocampus during early consolidation. A likely mechanism involves a transient redistribution of metabolic resources induced by HIIE, leading to a temporary downregulation of prefrontal activity ^51^. Since prefrontal engagement is crucial for memory reactivation, this temporary hypoactivity may have hindered the reinstatement of the spatial representation required to perform the SML accurately. Unlike current studies where HIIE had no effect on SML performance in accuracy and movement time assessed 24 hours later^52–54^, early impairments here in accuracy persisted at 24 hours. Performance recovered after a week, likely reflecting gradual refinement of the memory trace through repeated reactivation and reconsolidation^55^. The discrepancy with previous findings may be stem from task-specific factors, as the present bimanual 8-item sequence required more complex goal-based and motor coordination, likely engaging prefrontal substrates more strongly than simpler or implicit SML, thus increasing its susceptibility to early disruption. Although previous research has associated HIIE with reduced SICI and increased MEPs^22^, we found no changes in SICI excepted a difference between HIIE and REST independently of time and similar increases in MEPs in both HIIE and REST groups during early consolidation. Altogether, HIIE has been shown to enhance consolidation of visuomotor adaptation paradigms^17,50^, it may not confer similar benefits for explicit SML while HIIE promotes markers of plasticity (BDNF, lactate and MEP). In this context, moderate-intensity exercise may offer a more appropriate intervention strategy, as it could facilitate consolidation without overtaxing cognitive control resources critical for goal-based memory. This is supported by the findings of Rannaud Monany et al. (2023)^56^, who reported improved simpler SML consolidation when acquisition was followed by a moderate intensity exercise.

### Conclusion

Overall, this study provides novel evidence that a full night of SD selectively affects the movement time of motor sequence acquisition while preserving spatial accuracy, suggesting that the goal-based component of procedural memory remains resilient to acute sleep loss. Despite slower execution, accuracy was maintained and long-term consolidation was not compromised, suggesting that the core memory trace supporting SML can withstand acute disruption in sleep. These findings are particularly relevant given the prevalence of sleep disturbances in the general population, and they underscore the need to dissociate which components of memory are vulnerable versus preserved under sleep-deprived conditions. Looking ahead, future investigations should determine whether the robustness of goal-based motor memory observed under acute SD persists under chronic or ecologically valid forms of sleep restriction. Moreover, the unexpected decline in accuracy after HIIE, regardless of sleep status, challenges its presumed benefits as a post-learning enhancer. While further research is warranted to clarify the mechanisms involved, current results caution against the immediate use of HIIE following explicit SML.

## Author Contributions

**GD** contributed conceptualization, data curation, investigation, formal analysis, methodology, project administration, supervision, writing original draft, review & editing, validation; **TL** contributed conceptualization, resources, formal analysis, methodology, project administration, supervision, writing original draft, review & editing, validation; **JG** contributed investigation, methodology; **JB** contributed investigation, methodology; **VP** contributed conceptualization, resources, methodology, project administration, review & editing; **MP** contributed methodology, resources, project administration; **MP** contributed methodology, resources, project administration; **ES** contributed methodology, resources, project administration; **UD** contributed conceptualization, data curation, resources, formal analysis, methodology, project administration, supervision, writing original draft, review & editing, validation.

## Acknowledgments

The authors sincerely thank Dr. G. DEGLICOURT, Dr. L. BERTOLINO and Dr. G. CALLIES, as well as the entire department of functional respiratory testing of the Croix-Rousse hospital, for their invaluable support during participant inclusion and their assistance in handling blood samples. We warmly thank Ms. J. MAGAUD and Ms. L. BLANCO for their dedication, enthusiasm and valuable contributions to this work. We also thank the Clinical and Epidemiological Bases Unit (BCE) of the Public Health Division of the Hospices Civils de Lyon for their support in designing the eCRF and managing the study data.

## Data availability

The datasets used and/or analyzed during the current study are available from the corresponding author upon reasonable request.

## SUPPLEMENTAL INFORMATION

### Supplemental method

Experimental session for women was schedule during the first 10 days of the menstrual cycle, specifically during the follicular phase ^57^. At the end of the intervention (HIIE or REST), the subjective perception of both physical and cognitive workload were assessed via the National Aeronautics and Space Administration Task Load Index (NASA-TLX) questionnaire ^58^, as well as physical exhaustion with the rate of fatigue scale ^59^. Participants also completed the short version of the profile of mood state (POMS) questionnaire at the beginning of the experimental day and immediately after HIIE or REST.

#### Supplemental data

**Figure S1:**
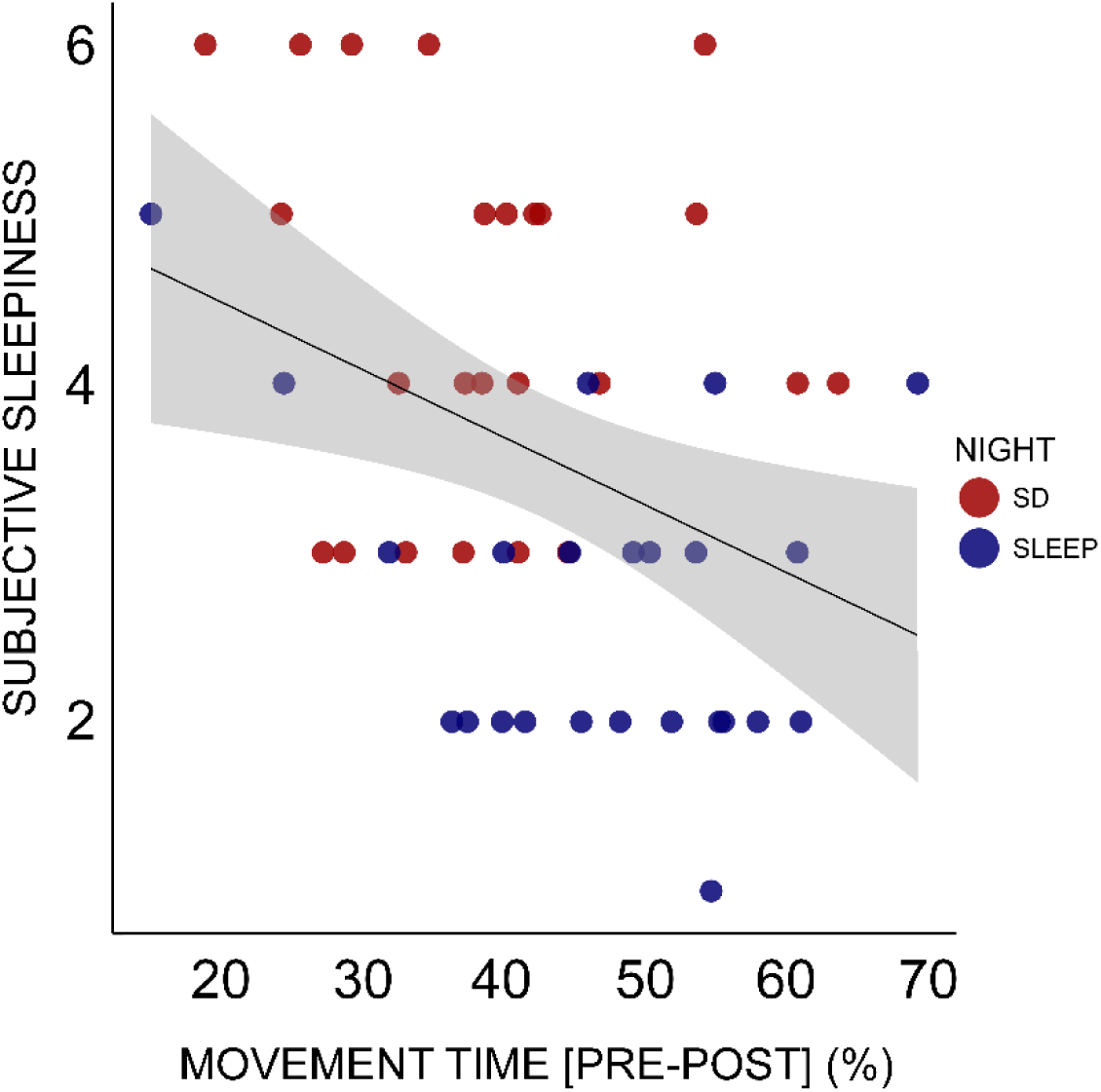
Negative correlation between subjective sleepiness and movement time.

**Figure S2:**
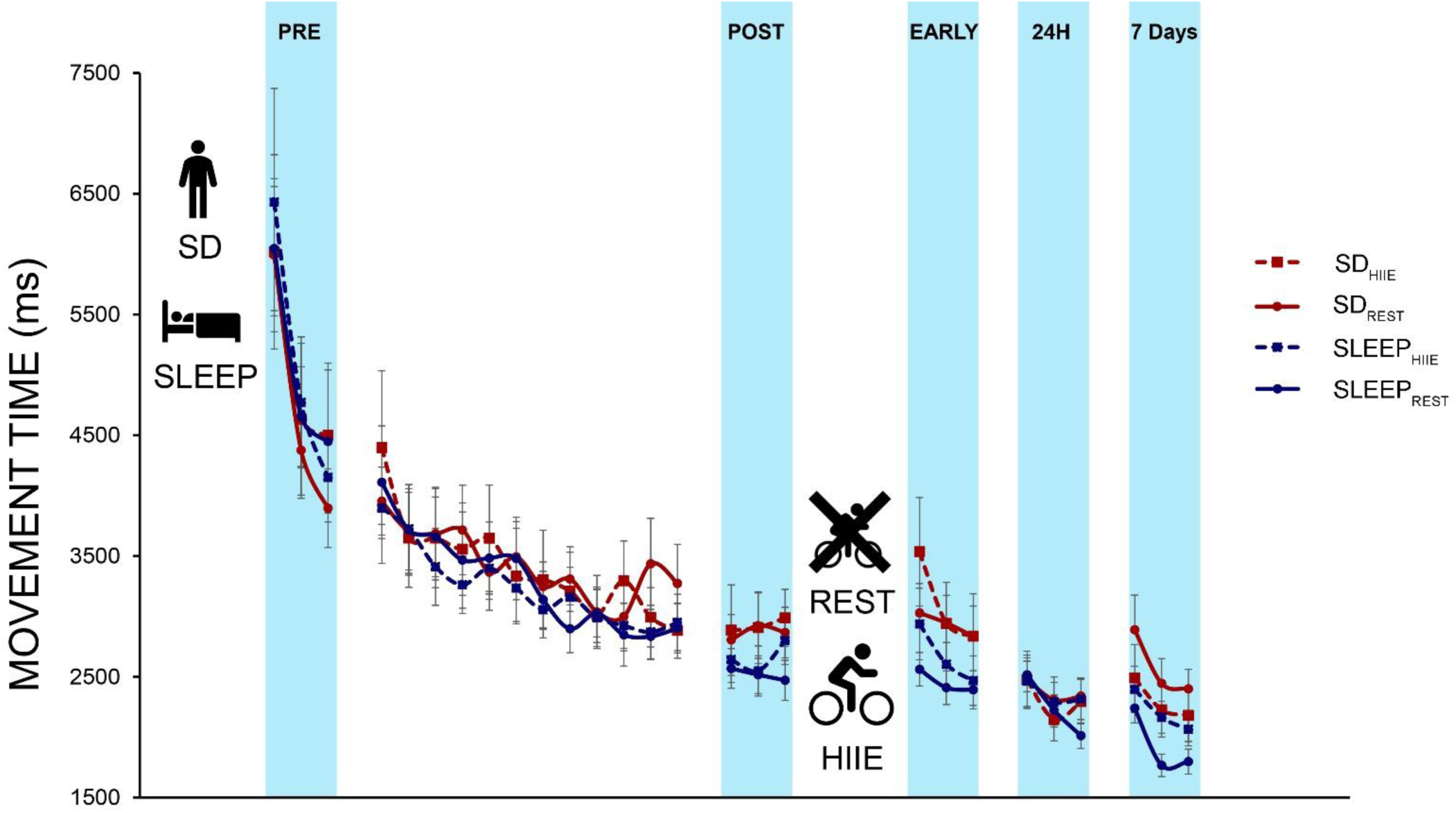
Performance curve for MOVEMENT TIME (ms). Errors bar represents the SEM.

**Table S1:**
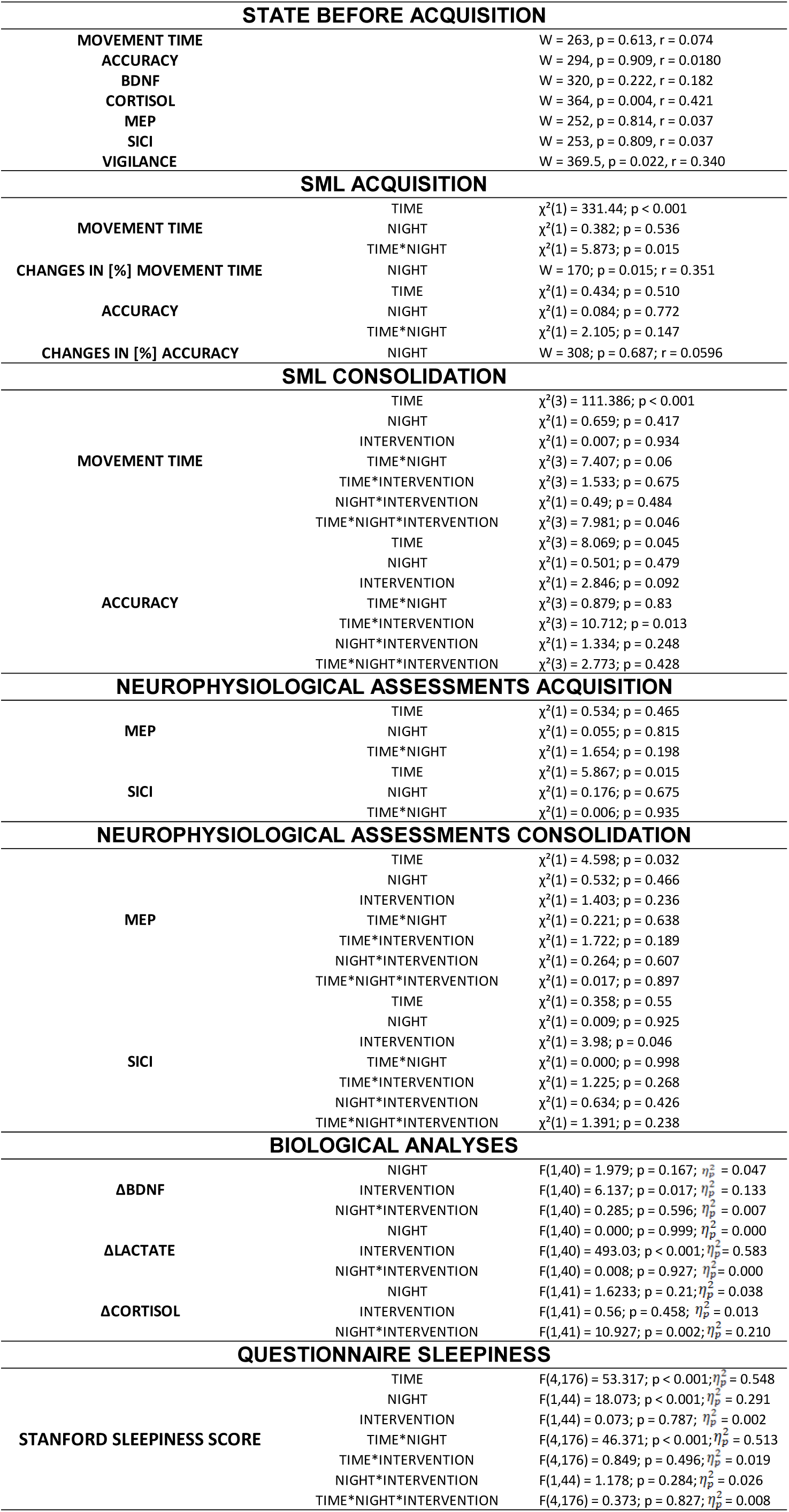
Detailed information of all analyses presented in the main manuscript.

## CONSOLIDATION

Changes in MOVEMENT TIME [%] across consolidation tests revealed no significant effects of NIGHT, INTERVENTION, or their interaction. From POST-ACQUISITION to EARLY-CONSOLIDATION no main effects of NIGHT (F(1,44) = 1.708; p = 0.198; 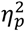 = 0.028), INTERVENTION (F(1,44) = 0.91; p = 0.345; 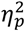 = 0.017), or NIGHT*INTERVENTION (F(1,44) = 0.083; p = 0.774; 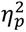 = 0.002) were observed. Similarly, from POST-ACQUISITION to 24 Hours, no significant effects emerged for NIGHT (F(1,44) = 1.836; p = 0.182; 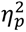 = 0.011), INTERVENTION (F(1,44) = 0.275; p = 0.602; 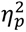 = 0.003), or their interaction (F(1,44) = 0.119; p = 0.732; 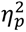 = 0.000). Finally, from POST-ACQUISITION to 7 Days, no main effect of NIGHT (F(1,44) = 0.120; p = 0.731; 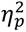 = 0.049), INTERVENTION (F(1,44) = 0.561; p = 0.458; 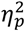 = 0.055), or NIGHT*INTERVENTION (F(1,44) = 2.289; p = 0.137; 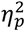 = 0.015) were found.

Changes in ACCURACY [%] revealed no significant effects of NIGHT, INTERVENTION, or their interaction across consolidation tests. From POST-ACQUISITION to EARLY-CONSOLIDATION, no effects were observed for NIGHT (F(1,44) = 0.005; p = 0.945; 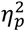 = 0.022), INTERVENTION (F(1,44) = 3.011; p = 0.09; 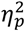 = 0.002), or interaction NIGHT*INTERVENTION (F(1,44) = 1.79; p = 0.188; 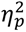 = 0.039). From POST-ACQUISITION to 24 Hours, no effects of NIGHT (F(1,43) = 1.086; p = 0.303; 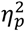 = 0.048), INTERVENTION (F(1,43) = 1.649; p = 0.206; 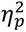 = 0.001), or interaction NIGHT*INTERVENTION (F(1,43) = 1.061; p = 0.309; 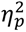 = 0.024) were detected. Likewise, from POST-ACQUISITION to 7 Days, no significant effects were found for NIGHT (F(1,44) = 0.239; p = 0.627; 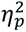 = 0.019), INTERVENTION (F(1,44) = 0.011; p = 0.916; 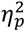 = 0.006), or their interaction (F(1,44) = 0.649; p = 0.425; 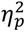 = 0.015).

